# Brain preparedness: The cortisol awakening response proacts dynamic organization of large-scale brain networks across emotional and executive functions

**DOI:** 10.1101/2024.02.22.581523

**Authors:** Yimeng Zeng, Bingsen Xiong, Hongyao Gao, Chao Liu, Changming Chen, Jianhui Wu, Shaozheng Qin

## Abstract

Emotion and cognition involve an intricate crosstalk of neural and endocrine systems that support allostatic processes for maintenance of dynamic equilibrium and rapid adaptation for upcoming challenges. As a hallmark of human endocrine activity, the cortisol awakening response (CAR) is recognized to play a critical role in modulating emotional and executive functions. Yet, the underlying mechanisms of such effects remain elusive. By leveraging pharmacological neuroimaging technique and Hidden Markov Modeling of brain state dynamics, we show that the CAR proactively modulates rapid reconfigurations (state) of large-scale brain networks across multi-task demands. Behaviorally, suppression of CAR proactively and selectively impaired accuracy for emotional discrimination task but not for working memory (WM). In parallel, suppressed CAR led to a decrease in the occurrence rate of brain state dominant to emotional processing, but an increase in brain state linking to executive control under high WM demand. Further energy-based analyses revealed an increase in transition frequency and sequence complexity along with an increased entropy during emotional tasks when suppressed CAR, suggesting a decreased energy supply. Moreover, an increased transition frequency was observed when shifting from neutral to emotional conditions, but an opposite pattern during WM task, with n decreased transition frequency shifts from low to high-executive demands. Our findings establish a causal link between CAR and dynamic allocation of neural resources for emotional and executive functions, suggesting a cognitive neuroendocrine account for CAR-mediated proactive effects and human allostasis.

## Introduction

For centuries, scientists have sought to unravel how the brain and endocrinal signals work in concert to support ever-changing cognitive and environmental demands. In theory, to sustain a dynamic equilibrium between internal milieu and external challenges, the interactions between endocrinal signals and brain systems actively engage neural resource allocation in an attempt to prospectively prepare for the upcoming challenges (McEwen et al. 2020, Bobba-Alves et al. 2022). Such active process has recently been conceptualized as ‘allostasis’ and is believed to substantially regulate human emotional and cognitive process, though the neurobiological mechanisms remain largely elusive. Of the endocrinal signals, the glucocorticoid stress hormone (cortisol in humans) plays a critical role in support of human emotion and cognition (Kunz-Ebrecht et al., 2004; Law et al., 2020; Schmidt et al., 2015). And the cortisol awakening response (CAR), in particular, as a robust rise within thirty to forty-five minutes after morning awakening, is thought to exhibit the proactive effects on human cognitive and emotional processes throughout the day (Clow et al., 2010; Xiong et al., 2021). As a hallmark of the hypothalamus-pituitary-adrenal (HPA) axis activity, the CAR is thus considered as a potential mediator of allostasis in humans (Clow et al., 2010). Although the effects of CAR on human cognition and emotion are widely documented at a behavioral level, our understanding of the underlying neurobiological mechanisms still remains in its infancy.

As a potential mediator of allostasis, the CAR should meet at least two following predictions according to the neurobiological models of allostasis: 1) responsible to modulate neural resource allocation for cognitive and affective processing in a proactive rather than reactive or passive manner, and 2) enable behavioral flexibility by dynamically reconfiguring functional brain networks in response to multiple task demands. Previous studies have well validated that the CAR is more than the mere release of cortisol with several unique features, including changes in attainment of consciousness and alertness (Clow et al. 2010), activating memory representations about future goal during stressful consequence (Fries et al. 2009), and modulating higher-order cognitive functions (Moriarty et al. 2014). As such, the CAR is postulated to exhibit an important preparation function for human brain and cognition. This preparation hypothesis is supported by the modulatory effect of CAR on core brain regions, including hippocampus, amygdala as well as prefrontal cortex (de Kloet et al. 2019), all of which possess abundant mineralocorticoid (MRs) and glucocorticoid receptors (GRs). By binding with these receptors, CAR could proactively modulate neural excitability in both local (Joëls et al. 1992) and regional connectivity among limbic structure and prefrontal regions (Xiong et al. 2021). However, how CAR could flexibly modulate multiple tasks processing (i.e. emotional and cognition) in a proactive manner remains elusive. Recent advances in network neuroscience have pointed out that dynamic (re)configurations of functional brain networks are essential to support human cognitive and behavioral flexibility (Bassett et al. 2006, Seeley et al. 2007, Bullmore et al. 2009, Vidaurre et al. 2017, Capouskova et al. 2022, Presigny et al. 2022). To date, however, little is known about whether and how CAR modulates dynamical organization of functional brain networks to support multiple cognitive and affective functions.

We believe the proactive and flexible effects of the CAR on human cognition and emotion could result from an interplay between CAR-initiated background tonic activity and stimulus-induced catecholaminergic phasic actions (van Stegeren et al. 2007, Joëls et al. 2009), both of which can act on large-scale neural networks via wide-spread neuromodulatory and endocrinal projections (Roozendaal et al. 2009). For emotional processing, the core hubs of salience network could be substantially modulated by both cortisol and stimulus-induced neuromodulators including norepinephrine (McReynolds et al. 2010, Menon et al. 2010, Young et al. 2017). Meanwhile, the interactions between cortisol and phasic dopaminergic signals that project to prefrontal cortex and several parietal, temporal regions could substantially modulate dynamic reconfigurations of executive control networks (Fornito et al. 2012, Ray et al. 2020, He et al. 2023). Together, the interplay between wide-spread cortisol and neuromodulator pertains to CAR’s ability to flexibly reconfigure network reconfigurations. However, few studies have directly investigated such CAR-related dynamic reconfigurations across brain networks. Recent advances in dynamical network modeling such as hidden Markov modeling (HMM) have been validated to probe time-resolved brain network configurations (or states), outperforming traditional sliding-window based machine learning approaches (Chang et al. 2010, Hutchison et al. 2013). Moreover, this approach can provide brain state dynamics including occurrence and transitions, thus ideal to probe CAR-related rapid configurations of brain networks under a multiple task context. Based on the neurobiological models of general cortisol in animal models and humans, we hypothesize that the CAR would dynamically regulate core network organization across emotional and executive processing.

Based on the allostasis theory, the CAR modulatory effects on cognitive and affective functions could further attribute to flexible reallocation of neurocognitive resources among large-scale functional brain networks. Such allocation of neural resource has been hypothesized that could efficiently regulate internal energy supply in an attempt to support specific neurocognitive process at the cost of others (Barulli et al. 2013, Hermans et al. 2014) and exhibit a tight link with system allostasis (McEwen 2000, McEwen et al. 2020). We believe such reallocation phenomenon could give mechanistic explanation regarding to why rapid reconfiguration could occur across distributed neurocognitive systems in a relatively short time period. Based on neurobiological studies (Xin et al. 2018, Shine 2019), neurotransmitters like norepinephrine as well as dopamine have been validated that could substantially affect local neural excitability and reshape the neural resource equilibrium in response to external changes, leading to higher neural and behavior adaption toward tasks (van Stegeren et al. 2007, Joëls et al. 2009). Notably, such neural resource allocation has been consistently linked to dynamical stability of brain systems (Hermans et al. 2014, Shine 2019, Munn et al. 2021). Thus, how to quantify CAR’s potential regulation toward dynamical system stability at a network level is essential to reveal reallocation of neural resource by CAR. Moreover, computational metrics like entropy and energy horizon have been proved to be feasible to quantify the dynamic reallocation between neurocognitive resources (Munn et al. 2021). According to dynamic organization of functional brain networks and above empirical observations, we thus hypothesize that the CAR would yield similar mechanisms to regulate system-level stability during affective and executive demands via flexibly modulating neural resource allocation across neurocognitive systems.

Here we test these hypotheses by combining a double blinded pharmacological manipulation (Cohort 1) dedicated to the CAR, in conjunction with a multi-task paradigm across three consecutive days. As shown in **Figure 1**, participants received either 0.5-mg dexamethasone (DXM) or placebo (vitamin C) on Day 1 night (20:00) to suppress CAR in the next morning. And saliva samples were collected at a total of 15 time points spanning over three consecutive days as a measure of cortisol level. Thereafter, participants underwent functional magnetic resonance imaging (fMRI) to perform a set of multiple tasks, including rest, emotion matching and working memory. The inclusion of resting state is to examine the effects of CAR on task-free intrinsic brain state dynamics as relative to task-dependent emotional and WM tasks implicated into the emotional salience network, and the frontoparietal executive network and default mode network. We first focused on the proactive effects of CAR on affective and executive control tasks. To further validate resource allocation hypothesis, we then implemented the HMM to probe dynamical organization of large-scale functional brain networks and investigate how CAR flexibly regulates brain state dynamics under a multi-task context. Finally, we developed information theory-based algorithms to compute shifts of system stability as well as entropy to examine whether CAR regulate neural resource reallocation across affective and cognitive tasks. Notably, we replicated most of the result in an independent cohort (Cohort 2, total 59 participants) and validate the generalizability and robustness of our results.

**Figure 1.**
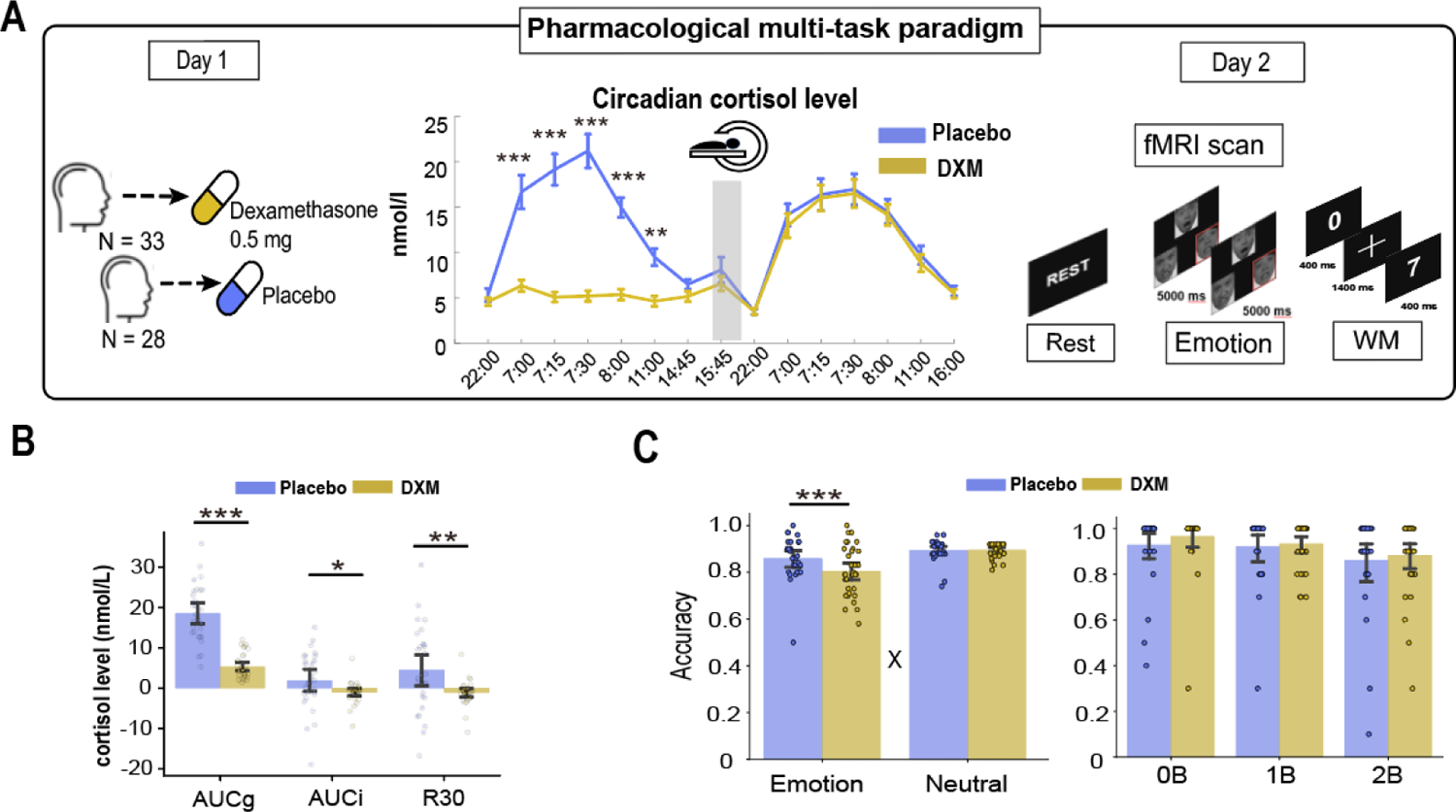
The CAR’s proactive effect on multi-task behavioral performance. **(A)** At first day night, participants were administered either 0.5 mg DXM or the same dosage placebo. On Day 2 afternoon, participants completed 3 consecutive tasks including rest, emotion matching and N-back. A total of time-stamped participants’ saliva samples was conducted throughout 3 days to measure CAR levels. **(B)** Bar graphs depict AUCg, AUCi and R30 for placebo and DXM groups, respectively. **(C)** Bar graphs depict accuracy of emotion matching task and N-back performance between task condition and groups. During emotion condition, placebo showed significantly higher accuracy compared with that in DXM group. Notes: *, p<0.05, **, p<0.01.

## 2. Methods

### 2.2 Participants

A total of 120 young, healthy, male college students were recruited in two cohorts, with 62 participants (mean age: 22.9 ±1.9; range: 20-24 years old) in Cohort 1 and 59 participants in Cohort 2 (mean age: 21.9 ±0.7; range: 18-27 years old). Given hormonal fluctuations during periodic menstrual cycle in female may impede a reliable assessment of the normative CAR, female participants were not included in the present study. All participants reported no history of neurological, psychiatric or endocrinal disorders. We first excluded participants if these factors not meet the requirements: tobacco or alcohol use, irregular sleep/wake rhythm, intense daily physical exercise, abnormal hearing or vision, predominant left-handedness, current periodontitis, stressful experience or major life events. More detail can be found in our previous study (Xiong et al. 2021). Informed written consent was obtained for each participant before the experiment and study protocol was approved by the Institutional Review Board for Human Participants at Beijing Normal University. The protocol with pharmacological manipulation was registered as a clinical trial before the experiment (https://register.clinicaltrials.gov/; Protocol ID: ICBIR_A_0098_002). Two participants in Cohort 1 were excluded from further analyses because of excessive head movement over 3 mm or averaged frame-wise displacement > 0.2mm during fMRI scanning according to the exclusion criteria by previous studies (Kottaram et al. 2019, Chen et al. 2022).

### 2.2 Pharmacological manipulation and cognitive tasks

For Cohort 1, participants were randomly assigned into the CAR-suppressed or control group. We employed the randomized, double-blind pharmacological manipulations in order to effectively suppress the cortisol awakening response (CAR). Participants in one group received a dose of 0.5 mg Dexamethasone (DXM group) and the control group received equal amount of Vitamin C (placebo group) at 20:00 on Day 1. Moreover, we implemented an independent Cohort 2 with the same multi-task design and subject’s natural CAR were measured.

In Cohort 1, to capture the diurnal cortisol responses as a function of time from morning to evening, we used a time-stamped dense sampling approach to collect a total of 15 saliva samples over 3 consecutive days (**Figure 1**): Day 1 at 22:00, Day 2 at 7:00 (awakening time), 7:15 (15 min after awakening), 7:30 (30 min after awakening), 8:00 (1 hour min after awakening), 11:00 before lunch, before and after fMRI scanning in the afternoon from 14:00 to 17:00 and 22:00 in the evening; Day 3 at 22:00; Day 2 at 7:00 (awakening time), 7:15 (15 min after awakening), 7:30 (30 min after awakening), 8:00 (1 hour min after awakening) and final 2 sampling at 11:00 and 16:00 respectively. Saliva samples were collected using Salivette collection device (Sarstedt, Germany). In Cohort 2, we collected morning cortisol samples within 1 hour immediately post-wakening from two consecutive days. Total 10 saliva samples were collected with 0, 15, 30 and 60 min after awakening on both Day 1 and Day 2. Extra 2 time points before and after fMRI scanning on Day 2.

We took several stringent procedures to assure that saliva was not affected by other factors not interested including 1) participants were asked not to brush teeth, drink or eat within 1 hour before sampling 2) participants were required to refrain from any alcohol, coffee, nicotine consumption and excessive exercise at least one day before experiment. Saliva samples were returned back to the laboratory and kept frozen (−20^。^c) until assay. After thawing and centrifuging at 3000 rpm for 5 min, the saliva samples were analyzed using an electrochemiluminescence immunoassay (ECLIA, Cobas e601, Roche Diagnostics, Mannheim, Germany) with sensitivity of 0.500 nmol/L. Intra and inter assay variations were below 10%. The CAR was computed by the area under the curve with respect to increase (AUCi) by the equation: AUCi = (S1+S2) x 15min/2 + (S2+S3) x 15min/2 + (S3+S4) x 30min/2 – S1 x (15 min +15 min + 30 min). The AUCi reflects the diurnal changes of CAR over time (Pruessner et al. 1997, Clow et al. 2010). Notably, we focus on CAR assessment mainly on Day 2, given our hypothesis and questions on the proactive role of the CAR on the brain state dynamics.

### 2.3 Multiple cognitive and affective tasks

At Day 2 afternoon, participants went through a consecutive multiple tasks involving Resting state, Emotion matching and N-back working memory. During rest, participants were instructed to open the eyes, relax and be still and scanning duration for 348s. During emotion matching task, participants performed a negatively emotional face-matching (vs. sensorimotor control) task. The task consisted of 10 blocks, each block containing 6 trios of images. Each block started with a cue for 5 s indicating either emotion or control condition, after which 6 trios of images were presented 5s each. For emotion condition block, participants need to select the emotion face from two candidate faces in the bottom of the screen that expressed the same emotion as the target expression on the top of the screen. For control condition, participants viewed the same number of trios of images and each block started with a cue for 5 s. Thereafter, participants need to select the scrambled shapes from two candidate shapes in the bottom of the screen that expressed the same shape as the target shape on the top of the screen. Notably, only negative facial expressions were included in the present study.

For N-back working memory task, it consisted of 12 blocks of alternating 0-,1- and 2-back conditions. Each block started with a 2-s cue indicating the experimental condition, followed by a pseudo-randomized sequence consisting of 15 digits. Each digit was presented for 400 ms followed by an inter-stimulus-interval of 1400 milliseconds. Blocks were interleaved by a jittered fixation ranging from 8 to 12 s, resulting in a mean inter-block duration of 38 s. During 0-back condition, participants need to detect whether the current digit was ‘1’. During the 1-back condition, participants were instructed to detect whether the current digit is the same with digit appeared 1 position back in the sequence. During the 2-back condition, participants were instructed to detect whether the current digit is the same with digit appeared 2 positions back in the sequence. Each sequence contained either 2 or 3 targets and participants were asked to make a button press with their right index finger as fast as possible when detecting a target. The employment of 3 distinct task for participants is to elicit sufficient and distinct cognitions covering most brain networks and better capture the dynamic shifts induced by cortisol awakening response. This multi-task procedure is proved to be efficient in eliciting distinct whole-brain network configurations in other studies focusing on network dynamicity (Telesford et al. 2016, Capouskova et al. 2022).

For Cohort 2, the task procedure is the same with Cohort 1, with consecutive tasks of rest, emotion matching and working memory, but during working memory we adopted a simplified design with only two WM loads (i.e., 0- and 2-back). Other parameters and task design are the same with Cohort 1.

### 2.4 Brain imaging data acquisition

Whole-brain images in both Cohorts were acquired on a Siemens 3.0T TRIO MRI scanner (Erlangen, Germany) in the National Key Laboratory of Cognitive Neuroscience and Learning & IDG/McGovern Institute for Brain Research at Beijing Normal University. Functional brain images were consecutively collected during resting state, emotion matching and N-back tasks, using gradient-recalled echo planar imaging (GR-EPI) sequence (axial slices = 33, volume repetition time = 2.0s, echo time = 30 ms, flip angle = 90, slice thickness = 4 mm, gap= 0.6mm, field of view = 200 x 200 mm, and voxel size = 3.1 x 3.1 x 4.6 mm). T1-weighted 3D magnetization-prepared rapid gradient echo sequence (slices = 192, volume repetition time = 2530ms, echo time = 3.45ms, flip angle = 7°, slice thickness =1 mm, field of view = 256 x 256 mm, voxel size = 1 x 1 x 1 mm^3^).

### 2.5 Functional data preprocessing and BOLD time series extraction

Image preprocessing was performed using Statistical Parametric Mapping (SPM12, http://www.fil.ion.ucl.ac.uk/spm). Preprocessing steps include 1) removing the first four volumes of functional images for rest, emotion matching and N-back task respectively, for signal equilibrium, 2) correcting for slice-timing acquisition, 3) realigned for head motion correction, 4) co-registered to the gray matter image segmented from subject’s own anatomical T1-weighted images, 5) spatially normalized into common stereotactic Montreal Neurological Institute (MNI) space, and 6) images were finally resampled into 2-mm isotropic voxels and smoothed by isotropic 3-D Gaussian kernel with 6 mm full-width at half-maximum. We employed a pre-defined whole brain network template (Allen et al., 2014). This set of templates yield a total of 14 networks that covering most cortical and subcortical regions. We next extracted pre-processed BOLD time series for each task by using the 14 network templates, which results in a time series matrix (14 brain networks) x (task durations). For Cohort 1, time series matrix for rest is 14-by-170, for emotional task is 14-by-150 and for N-back task is 14-by-228. For Cohort 2, time series matrix for rest is 14×176 for emotional task is 14-by-150 and for N-back is 14-by-228. For both Cohorts, the white matter and cerebrospinal fluid, as well as 24-head movement signals were further used as regressor to remove out from time series.

Notably, we only removed the low frequency drift with high-pass filtering at 0.08 Hz for time series from 3 tasks to keep consistent interpretability between rest and task in later modeling and analysis. Preprocessed time series of 14 networks for the three tasks were temporally concatenated together for each participant, yielding a 14-by-548 time series matrix for Cohort 1 and 14-by-554 time series matrix for Cohort 2. The concatenated time series were normalized per column to yield zero mean and standard deviation. Finally, normalized time series matrices for each participant were combined to yield a N-by-14-T participants matrix ready for the HMM modeling --- where N and T represent the number of participants and the length of time series for three tasks.

### 2.6 Hidden Markov Model for multiple tasks

The HMM considers a set of multidimensional input signals that can be modelled as a sequence of latent discrete states occurring and omitting as time evolves. Each state is defined as both mean of each dimension and covariance among each dimension, generated by the Gaussian distribution. Here we consider that the human brain dynamically reconfigures distinct brain networks to meet ever-changing requirement of external cognitive and affective tasks. As such, we employed a set of 14 canonical brain networks derived by (Shirer et al. 2012) as pre-defined ROIs to cover potential network interactions during multiple tasks. These ROIs have been widely used by previous studies focusing on large-scale functional brain networks (Kottaram et al. 2019, Meer et al. 2020). After obtaining concatenated time series matrices, we employed a HMM-MAR toolbox (https://github.com/OHBA-analysis/HMM-MAR) to estimate potential dynamics underlying the data. Akin to previous studies (Vidaurre et al. 2017, Stevner et al. 2019, Meer et al. 2020), we chose a number of 10 states to capture various brain state dynamics covering three different tasks. We also searched parameter ranging higher than 10 states and found exist inefficient state which rarely occurred across the whole task conditions. The HMM model yielded three major output metrics: 1) Decoded state sequence for each time point, each subject, based on Viterbi path decoding; 2) Each state’s mean and covariance matrices derived from corresponding estimated mean across 14 brain networks (**Fig. 2A**), as well as fractional occupancy (relative occurrence rate of specific state in different task conditions or blocks); 3) Transition matrices among 10 distinct states, calculated by the likelihood of switching between pair of specified states given task conditions or blocks. Thus, we could calculate subject-specific and task-condition dependent transition behavior for later comparison between groups.

**Figure 2.**
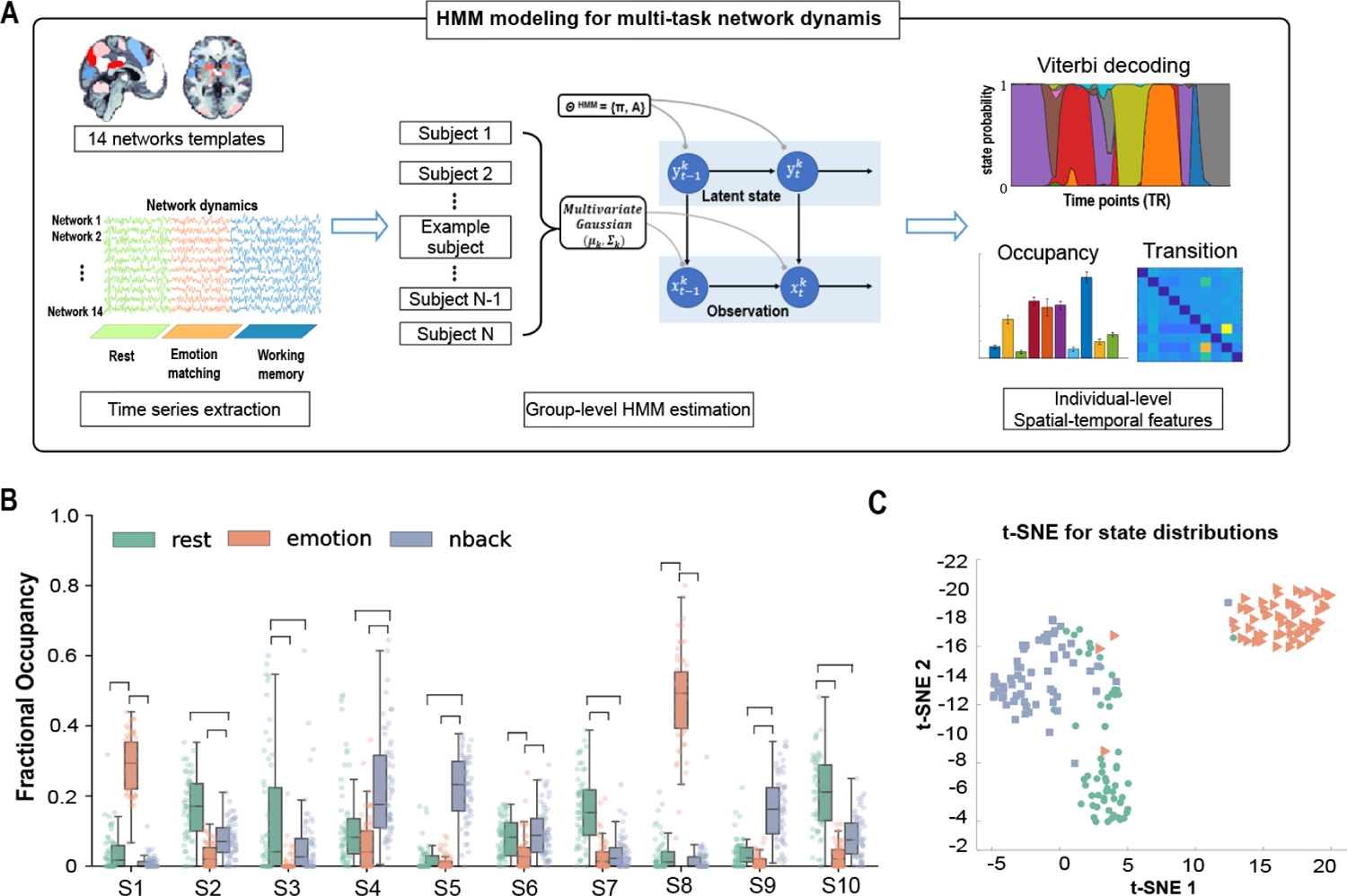
Spatiotemporal patterns of each brain state under multiple cognitive and affective tasks. **(A)** BOLD time series were extracted from 14 prior-defined brain networks of interest. The Hidden Markov models were implemented to estimate network dynamics throughout whole experiment. (**B)** Relative occupancy rate of each brain state during rest, emotion condition and N-back. Horizontal lines represent significant differences in fractional occupancy (p < 0.001, FDR corrected) among three task conditions for each state. (**C)** t-SNE embedding for distribution of occupancy rates of three tasks. Notes: *p_corr_<0.05, **p_corr_<0.01, ***p_corr_<0.001

### 2.7 t-SNE embedding visualization

To capture the distribution structure under distinct task conditions, we utilized a t-distributed stochastic neighbor embedding (t-SNE) visualization technique(Van der Maaten et al. 2008), which can represent high-dimensional data into a reduced dimension while preserving the local distances between data points. In our study, we defined states distribution under task condition as a 10-element vector, each element in the vector represents relative fractional occupancy of brain states per task condition. As such, we created the distribution vectors for all 3 conditions for each subject, and we then employed t-SNE analysis to plot the distribution vector across conditions per participant and visualize the relative distance among three conditions.

### 2.8 Knee point analysis

A knee point analysis based on previous study (Bijanzadeh et al. 2022) was employed to determine the dominate states for each condition. We first ranked average fractional occupancy of all brain states for rest, emotion matching and n-back tasks separately. Then, we calculated accumulative fractional occupancy curve and employed ‘kneedle’ algorithm (Satopaa et al. 2011) to detect knee point, defined by a significant transient point in the maximum curvature, as a threshold to select the most dominated state for each task. We conducted multiple comparison correction procedures based on the dominate states detected by knee point analysis for subsequent statistical analyses.

### 2.9 Neurosynth reversed inference

To reveal the potential neurocognitive processes associated with each state feature, we took advantage of meta-analysis using reverse-inferred similarity-based algorithm by the CANlab toolbox (https://canlab.github.io/). Consistent with previous studies (Yarkoni et al. 2011, Meer et al. 2020), we correlated each state’s activation pattern with the topic term-derived maps from the large-scale Neurosynth platform including 11406 studies. After obtaining similarity for the topics involved for each state, we chose top 10 topics with the highest positive similarity in candidate topics for each state as reverse-inferred cognition process underlying each brain state (only including topics that have positive similarity). We visualized each state’s related terms in a word-cloud plot and each topic’s size is based on its relative similarity value in that state.

### 2.10 Entropy analysis

To infer the relative energy level, we computed the entropy as a representative index of system energy. Based on the informational theory, entropy measures the uncertainty of a stochastic system and the increased entropy can be associated with reduced system’s uncertainty, pointing to an increase in system energy (Ezaki et al. 2017, Munn et al. 2021). In the present study, we calculated the entropy based on the probability of each state’s occupancy by the following formula:

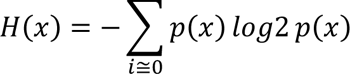

Where p(x) is the probability of each brain state’s occupancy.

### 2.11 Community detection

To detect potential transition modules underlying state evolution, we first calculated a group-wise transition matrix derived from the HMM models, where i and j represent transitions from state i to state j. Then, we employed Louvian community detection algorithm to obtain modules within transition matrix for each subject per task condition.

### 2.12 Transition centrality for particular state

To measure to what extent one given state is likely to maintain within itself from a transition perspective, we defined an index called transition centrality. First, we defined the overall transitions that other states transition toward specific state as T_toward,i_ and the overall transitions such specific state transit to other states T_away,i_. We refer these two metrics as toward transition strength and away transition strength. We calculated these two indexes based on the estimated transition matrix from the HMM models as follows:

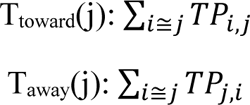

Where TP is the transition matrix and *TP*_i,j_ is the transition probability of state I at the time point of t-1 to state j at t. Finally, the transition centrality is defined as:

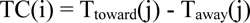

### 2.14 Transition frequency and complexity of state sequence

To measure the overall transition frequency in a given state sequence, we calculated transition frequency index to examine transition behavior under specific task condition (Cornblath et al. 2020). That is, given a state sequence under one task condition, we quantified all transitions between paired brain states while excluding self-transition. Assuming that we have a state sequence: D(condition_ij_) = S_4_S_7_S_7_S_7_S_7_S_7_S_7_S_4_S_4_S_4_S_4_. Its transition frequency is 2(transition behavior)/12(total length) equals to 1/6.

To test the robustness of our result for transition flexibility difference between groups, we implemented another algorithm called ‘substate sequences complexity’ (Crutchfield 2012, Clawson et al. 2019). This algorithm defines any state sequences as symbolic streams of words and measure the complexity of a given state sequences based on to what extent the list can be compressed into a shorter version of sequence without losing information of that sequence. This algorithm conceives that every meaningful sequence in the world should have compressed information between total randomness (which has the highest complexity since it is by no means feasible to compress) and total regular (which has the lowest complexity since it can be represented by few compressed words). In our study, we can also conceive state transition sequence as a stream of symbolic letters and use the same algorithm to measure the complexity of transition sequence. For instance, assuming that we have a state sequence D(t) = S_4_S_7_S_7_S_7_S_7_S_7_S_7_S_4_S_4_S_4_S_4_ with 10 letters. Based on complexity calculation, this sequence can be compressed as S_4_ 1 S_7_ 5 S_4_ 4 with 6 words. We thus can determine that the compression ratio of the given sequence is 6/10 equal to 0.5. We used this algorithm to test whether the complexity between placebo and DXM groups is still differed from each other, as an alternative metric to measure the underlying transition dynamics between two groups.

## 3. Results

### 3.1 Effectiveness of CAR suppression and its effects on behavioral performance

We first verified the effectiveness of CAR suppression and its proactive effects on multi-task behavioral performance (**Fig. 1A**). At Day 1 night, participants received either 0.5mg Dexamethasone or placebo (Vitamin C) to suppress the CAR in next morning. After that, a total of 15 saliva samplings were gathered from Day 1 at 22:00 through Day 2 into Day 3. At Day 2 afternoon, participants underwent fMRI scanning while they were performing 3 consecutive tasks. We first compared cortisol level between two groups from Day 1 to Day 3 by conducting a 2 (Group: DXM and placebo)-by-16 (Time: 15 samples) ANOVA analysis. Results revealed a significant main effect of Group (*F_1,57_* = 16.78, *p* < 0.001) and Group-by-Time interaction effect (*F_14,798_* = 19.91, *p* < 0.001). No significant group difference in cortisol levels were observed before and after fMRI scanning on Day 2 (All *ps* > 0.23). We further investigated group differences in the CAR based on three metrics, including AUCi, AUCg and R30, respectively (**Fig. 1B**). Two sample-t tests revealed significantly lower AUCg (*t_59_*=-9.61, *p* < 10^-13^), AUCi (*t_59_* = −2.10, *p* = 0.04) and R30 (*t_59_* = −2.93, *p* = 0.005) in the DXM group than Placebo controls, further validating the effectiveness of CAR suppression.

We then investigated how the CAR influences subsequent behavioral performance in emotion matching and working memory. For emotion matching, we conducted separate repeated-measure analysis of variance (ANOVA) for accuracy, with emotion versus neutral as within-subject factor and Group (DXM and Placebo) as between-subject factor. As shown in **Fig. 1C**, this revealed significant main effects of both group (*F_1,59_* = 2.94, *p* = 0.092) and emotion condition (*F_1,59_* = 27.45, *p* <0.001), as well as interaction effect (*F_1,59_* = 5.65, *p* = 0.021).

Post-hoc t-tests revealed higher accuracy of emotion condition in placebo group compared with that in DXM group (*t_59_* = 2.13, *p* = 0.037), but no significant difference in neutral condition (*t_59_* = −0.14, *p* = 0.89). Parallel analysis for accuracy data in working memory task revealed significant main effect of WM load (*F_1,59_* = 9.81, *p* = 0.003), but not interaction effect (*F_1,59_* = 0.12, *p* = 0.73). Additionally, we also analyzed efficiency measure accuracy divided by response time as another metric to quantify subject’s performance and found similar results as CAR selectively support performance of emotion matching (**Supplement Fig. 1**). These results indicate the effectiveness of CAR suppression by our pharmacological manipulation and a selective reduction in behavioral performance during emotional discrimination task.

### 3.2 Proactive effects of the CAR on brain state dynamics across multiple tasks

To detect brain state dynamics during multiple tasks, we implemented the HMM model for BOLD time series of 14 brain networks during resting state, emotional and working memory tasks collapsing across two groups (Fig. 2A). Model output 10 inferred latent brain states with distinct spatial features each across three tasks (**Supplement Fig.2**). We first compared relative fractional occupancy of each state under rest, emotional and working memory respectively. We found distinct state occurrence distributions among three tasks (Fig. 2B), with several brain states showing significantly higher fractional occupancy under specific tasks. We also calculated mean life time of each state during different tasks, and found distinct duration patterns under each task (**Supplement Fig. 3**). From a holistic view, we further implemented t-SNE algorithm to visualize the distribution of brain states under each task condition. This analysis revealed a low-dimensional clustering phenomenon showing distinct state distributions among each task (Fig. 2C). Notably, we conducted similar analysis for Cohort 2 and found a highly similar pattern of results (**Supplement Fig. 4**).

Next, we investigated how the CAR modulates brain state dynamics during multiple task context by comparing fractional occupancy under each task between groups. First, we conducted a knee-point analysis to identify brain states that most common occurred under each task (Fig. 3A). This revealed 4 dominate states in rest, whereas 2 and 3 dominate states for emotion matching and N-back respectively. (Fig. 3B**-D**). We therefore focused on these dominate state under each task in subsequent analyses. Two sample t tests for emotional task revealed significantly higher occupancy rate of core-emotional state (State 8) in the placebo than DXM group (t_303_ = 2.39, p = 0.03, FDR corrected), no difference in the neutral condition (Fig. 3C). Parallel analysis for N-back task (Fig. 3D) revealed significantly higher occupancy rate of low-executive state (State 2) during 1-back conditions in the placebo than DXM group (*t_242_* = 3.13, *p* < 0.01, FDR corrected), but marginally significant during 2-back condition (*t_242_* = 2.32, *p* = 0.06, FDR corrected). Interestingly, there is no reliable group difference during rest dominate state (all *ps* > 0.79) (Fig. 3B).

**Figure 3.**
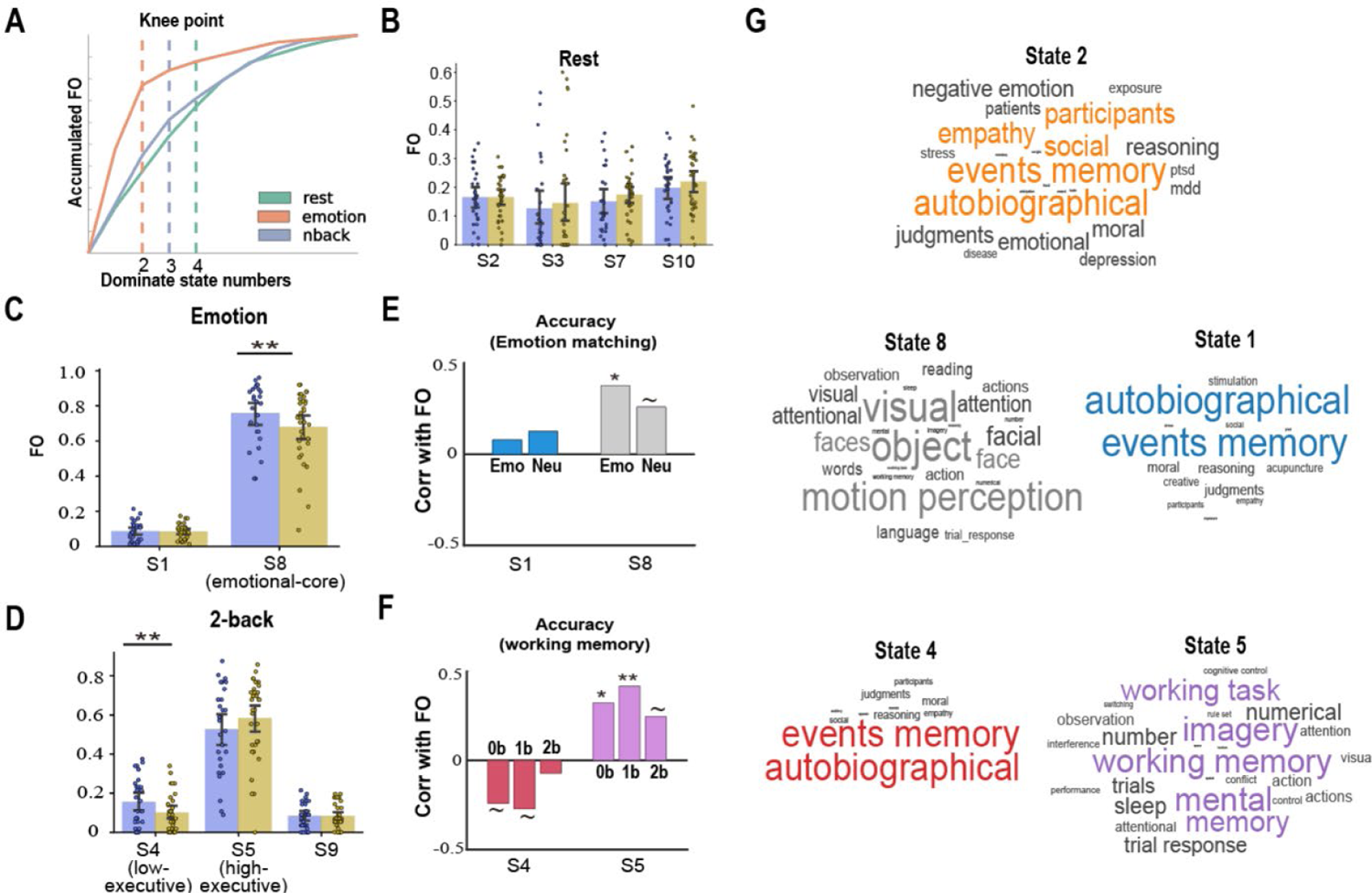
Distinct occupancy rates of core brain states showing significant differences between placebo and DXM groups. **(A)** Dominate state selection based on knee point analysis, vertical dashed lines represent corresponding dominate state numbers determined by knee point for each task separately. **(B)** Fractional occupancy difference between placebo and DXM group under resting and emotion matching **(C)** as well as N-back **(D)**. Error bar depict standard error of the mean. **(E-F)**. Correlations between fractional occupancy of dominate state and corresponding performance during emotion matching and N-back tasks. **(G)** Word-cloud plots for topic-based decoding of state properties using NeuroSynth. The font size is based on similarity value between activity map of each brain state and Neurosynth topics for each decoding. Plot colors is consistent with pre-defined color of each state. Notes: *p_corr_<0.05, **p_corr_<0.01, ***p_corr_<0.001.

To examine whether above group differences in brain state metrics are associated with task performance, we conducted several correlational analyses by collapsing across DXM and placebo groups. For emotional task, we found a marginally positive correlation between fractional occupancy of core emotional state and accuracy in emotional discrimination (r =0.38, p = 0.01, FDR corrected) (Fig. 3E). For N-back task, however, we found a negative correlation between fractional occupancy of low-executive control state and accuracy in 0-back (*r* =-0.24, *p* < 0.001) and 1-back (*r* =-0.27, *p* < 0.01) (all FDR corrected). Notably, the high executive control state (State 5) is positively correlated with accuracy in with 0-back (*r* = 0.32, *p* = 0.03), 1-back (*r* = 0.42, *p* < 0.01) and 2-back (*r* = 0.25, *p* = 0.07) (all FDR corrected) (Fig. 3F). To further characterize these states, we employed meta-analytic decoding approach to assess the similarity of each brain state’s network activation maps with the Neurosynth-based topic database (see Methods). As illustrated in Fig. 3G, three major brain states showing prominent group differences can be classified into following systems: 1) core emotional state linking to the topics of visual and face processing critical for emotion discrimination, 2) low executive-control state linking to the topics of autobiography, consistent with its relative low activity across all 14 networks, and 3) high executive-control state closely linked to working memory and numerical processing in our study. Again, in Cohort 2 we also identified a positive correlation between core emotional brain state and the CAR (R30), and occupancy rate of this affective state (*r* = 0.29, permuted *p* = 0.03) and exhibited a positive correlation with emotional performance (**Supplement Fig. 6C**).

In sum, these results indicate that the CAR proactively promoted occupancy rate of core emotional state during emotion task, while increased FO for low executive-control state during high-load working memory task.

### 3.3 Proactive effects of the CAR on brain state transitions cross multiple task contexts

Furthermore, the structure of transitions represents dynamical reconfigurations among brain states and we investigated how the CAR proactively modulates brain state transitions during multiple cognitive and affective task contexts. We first derived transition probability metrics among brain states based on each participant’s Viterbi decoding path and calculated the transition matric during each task condition (Methods). Louvian community detection (edge inclusion threshold: 0.2) revealed three communities showing highest within module connectivity, and the structure of community is highly consistent with the structure of dominate states based on previous knee analysis (Fig. 4A). Notably, we implemented different threshold and found robust community clustering results (**Supplement Fig. 5**). We also compared transition difference path-by-path between tasks, and found significantly higher transition probability within dominant brain states under each task conditions (non-parametric permutation: all *ps* < 0.001) (Fig. 4B). More importantly, we investigated whether the CAR modulates transition behaviors of core-emotional state and core executive state dominated in emotional and working memory conditions respectively. To achieve this, we quantified each state’s transition centrality. As shown in Fig. 4C, for core-emotional state we found marginally higher transition centrality in placebo than DXM group (permuted *p* = 0.061). Mixed-effect repeated measure ANOVA for 2-back condition revealed marginally significant interaction effect for transition centrality of executive-inactive and executive-active (*F_1,242_* = 3.15, *p* = 0.077). Post-hoc t-tests found significant higher transition centrality of executive-inactive state (permuted *p* = 0.015), but an opposite pattern for executive-active state for the placebo than DXM group (permuted *p* = 0.036).

**Figure 4.**
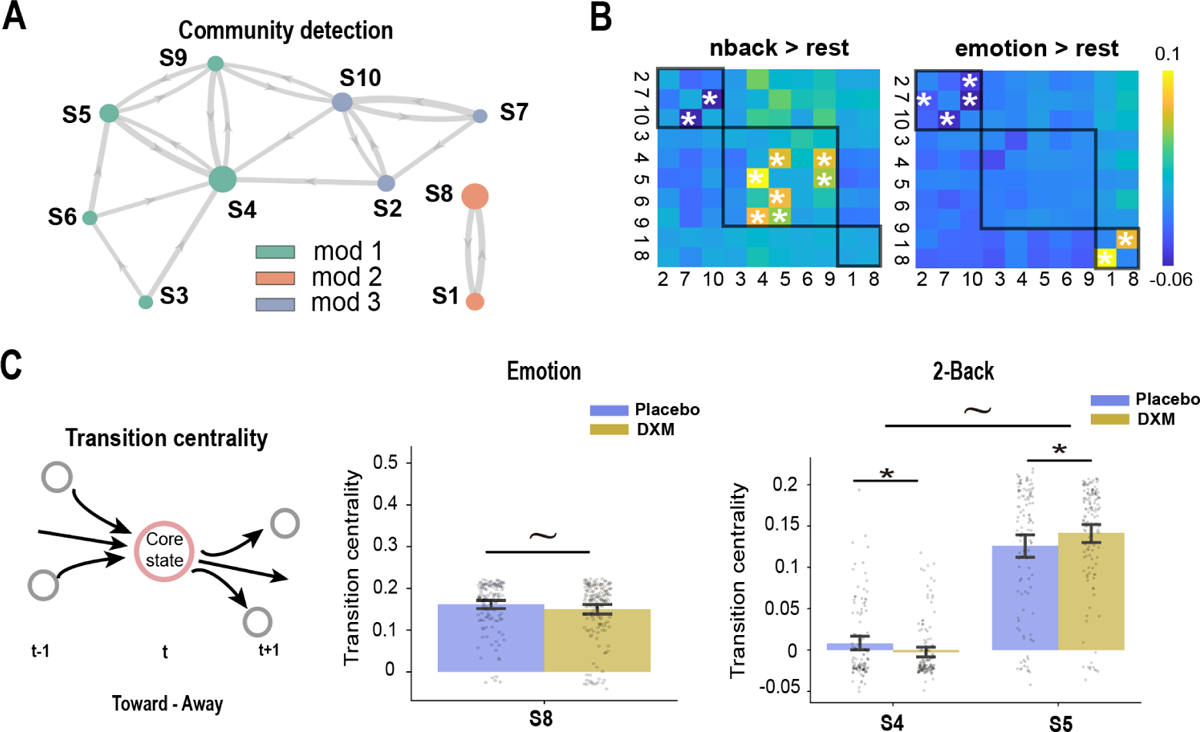
Differences in brain state transitions between placebo and DXM groups. **(A)** Louvian-detected community modules based on transition probabilities among all states. Colored background indicates states belonging to pre-defined dominate states for each condition. **(B)** Transition difference between different tasks. States were re-ordered based on their community assignment. **(C)** Transition centrality for affective-active state in emotion condition. **(D)** Transition centrality for dominate executive-inactive and executive-active in 2-back condition. Notes: ∼p<0.10, *p<0.05, **p<0.01.

Finally, we investigated whether the CAR influences the neurocognitive resource allocation representing an efficient way to regulate cognitive and affective processing. Based on empirical observations and assumptions on functional organization of large-scale brain networks(Ezaki et al. 2017, Munn et al. 2021), we speculate that higher allocation of neurocognitive resource can substantially increase the system-level stability of brain state dynamics. Thus, we first calculated overall transition frequency during each task condition as a way to quantify such system stability (Fig. 5A). We found significantly lower transition frequency during emotional condition in the placebo than DXM group (permuted *p* = 0.009). However, we did not find any group difference for working memory condition (permuted *p* = 0.3). Notably, we employed a sequence complexity analysis as an alternative metric of system stability (Fig. 5B). Again, we found marginally significantly lower complexity during emotional condition in the placebo than DXM group (permuted *p* = 0.068), while no difference during 2-back (permuted *p* = 0.52). We also implemented an entropy analysis based on state occupancy distribution as another measurement of resource allocation. Two sample t test revealed significantly lower entropy (i.e., higher system stability) in the emotion condition for the placebo than DXM group (*t_303_*: −2.31, *p* = 0.02). This is reminiscent of above results based on transition frequency or complexity analysis.

**Figure 5.**
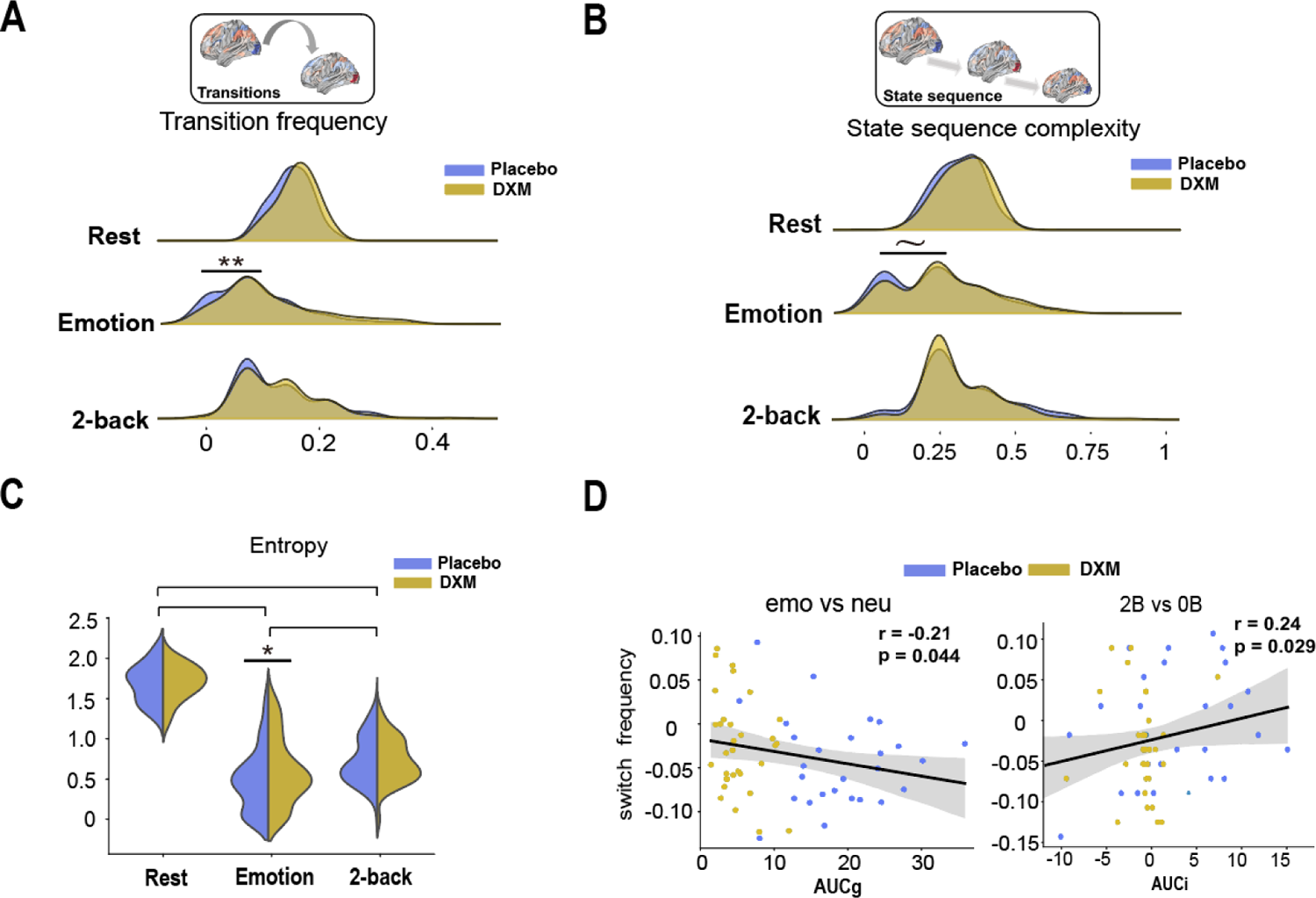
Transition dynamic differences between placebo and DXM groups. **(A)** Switch frequency among rest, emotional and 2-back condition. Switching frequency is measured by how frequent different states switch. **(B)** State sequence complexity among rest, emotion and 2-back condition. State sequence complexity is measured by the compressibility of a given state sequence. **(C)** Switching frequency shifts during emotional and N-back is correlated with cortisol awakening response, respectively. **(D)** Entropy value for rest, emotion and 2-back between DXM and placebo groups. Values are calculated based on its relative state occupancy rate distributions. Horizontal lines represent significant between group difference in entropy. Notes: ∼p<0.10, *p<0.05, **p<0.01.

Furthermore, we found that individual differences in the CAR level were correlated with their brain dynamic transitions. As shown in Fig. 5D, participant’s AUCg was significantly negatively correlated with transition frequency shift from neutral to emotion conditions (*r* = −0.21, permuted *p* = 0.043). Meanwhile, we fond significant negative correlation between subject’s AUCi is significantly positively correlated with transition frequency shift from 0-back to 2-back conditions (*r* = −0.24, permuted *p* **=** 0.029). Notably, we found a similar relationship between switching frequency and individual’s CAR (**Supplement** Fig. 6). To be specific, we found individual’s switching frequency shift (emotion vs neutral) was negatively correlated with corresponding AUCg (*r* = −0.23, permuted *p* **=** 0.035), but an opposite correlation between individual’s switching frequency shift (2-back vs 0-back) and corresponding AUCg (*r* = 0.25, permuted *p* = 0.026). Together, these results indicate that the CAR proactively modulates energy allocation reflecting by brain state dynamics under multiple task context.

## 4. Discussion

By leveraging pharmacological manipulations and cognitive neuroimaging in a multi-task paradigm, our study investigates how the CAR, as a candidate of allostasis, modulates rapid reconfiguration patterns of functional brain networks under executive and emotional demands. Behaviorally, CAR in the morning could selectively and proactively improve discrimination ability of negative facial expressions but not work memory performance later in the afternoon after the same day. Further Hidden Markov modeling revealed that such selective modulation on emotional processing was supported by dynamic reconfigurations of brain networks during transient tasks, with enhanced core emotion brain state dynamics but impaired executive-related brain dynamics during affective and executive processing, respectively. Further analyses of brain state stability and transitions revealed an increase in energy supply for emotional processing but an opposite pattern during WM, suggesting a flexible allocation of neural resources to support affective and executive tasks. Finally, we replicated our major results in an independent Cohort 2. Collectively, our findings establish a causal link of CAR in the morning and its proactive modulation on affective and executive functions, most likely through rapid reconfiguration of large-scale functional brain networks to enable flexible reallocation of neurocognitive resource in a multiple task context. These findings will be discussed under a framework of CAR serving as human allostasis.

### CAR proactively and selectively promotes emotional but not executive processing through dynamic organization of dominate brain states

Behaviorally, we observed a selective modulation of CAR linking to improved accuracy for discriminating negatively facial expressions, but not for working memory performance. Based on the CAR-mediated preparation hypothesis (Kunz-Ebrecht et al. 2004, Steptoe et al. 2016, Xiong et al. 2021), CAR can proactively initiate a tonic modulation on subsequent neuroendocrinal and behavioral adaption even several hours after its first burst in the morning, most likely via relatively slow genomic actions (Arnsten 2009). More importantly, such background activity could further interplay with neurotransmitters involving norepinephrine and dopamine when facing with affective and/or cognitive demands (namely phasic effect) (van Stegeren et al. 2007, Joëls et al. 2009, Munn et al. 2021, Xiong et al. 2021). Thus, we speculate that the tonic effect of CAR could more likely interact with task-dependent activity of neuromodulators to flexibly regulate emotional and executive process via dynamically regulating network reorganization.

Indeed, our dynamic network modeling (HMM) revealed that the CAR proactively increased the occurrence and transition centrality of brain state dominant to emotional processing (‘emotional brain state’), characterized by higher engagement of regions in the salience network along with posterior visual system. Notably, the presence of such emotional state has been found to promote goal-directed visual attention for the upcoming affective stimuli (Hermans et al. 2014), in line with our observed positive correlation between occurrence of this emotional brain state and individual difference in emotional performance. Based on previous observations linking CAR with negative affect on a behavioral level (Clow et al. 2010, Law et al. 2020), such enhanced occurrence of core emotional mode could attribute to the interplay between tonic effect of CAR and noradrenaline released by locus coeruleus (LC) targeting to the core hubs of emotional salience network including amygdala, which yields both β1-adrenoceptors and dopaminergic specific D1-receptors (Lamont et al. 1998, Birnbaum et al. 1999). By doing so, CAR could augment the occurrence of emotional brain state that optimal for affective-related processing and support affective functioning.

In contrast, during high-load working memory the suppressed CAR is correlated with higher occurrence and transition centrality of the ‘low executive state’, characterized by generally lower activity of all brain networks as well as a decreased transition centrality of ‘high executive state’, featured by high engagement of the executive network. These results are in line with previous behavior studies linking CAR with impaired executive functions(Devilbiss et al. 2012, Moriarty et al. 2014). Predictions of the neurobiological models from rodent and monkeys have posited that CAR-mediated tonic effect (corticosteroids) works in concert with task-driven phasic noradrenergic actions (catecholamines as well as norepinephrine) could orchestrate to influence executive networks, possibly through the low-affinity of α-1-adrenoceptors and β1-adrenoceptors in PFC (Birnbaum et al. 1999, Arnsten 2009, Hermans et al. 2014), leading to a decreased ability to maintain optimal executive brain states. And the fact that occurrence rate of low and high executive state is oppositely correlated with individual working memory performance further support our statement.

Taken together, our observed proactive effects of the CAR on emotional and executive functions most likely result from an interplay between CAR-mediated tonic background activity and task-invoked phasic catecholaminergic activity acting on dynamic reconfigurations of functional brain networks across emotional and executive brain states. Such dynamical regulation on the network-level substantially extends previous studies linking CAR to local activation and connectivity in a single task paradigm (van Stegeren et al. 2007, Xiong et al. 2021).

### CAR modulates neural resource allocation during emotional and executive processing

Our observed selective enhancement of system stability during emotional processing further suggests a dynamic shift in resource allocation across emotional and executive neurocognitive systems. In theory, such allocation of neural resource could substantially regulate the energy landscape of brain systems and determine whether brain could reach optimal mode in response to external stimuli (Gu et al. 2017, Munn et al. 2021, Capouskova et al. 2022). And several studies have emphasized that the endocrinal and neuro-modulatory systems could substantially modulate large-scale brain activity, possibly by modulating the given energy supply that belong to each neurocognitive system (Hermans et al. 2014, Xin et al. 2018, Munn et al. 2021).

Beyond the conventional approaches, by quantifying system dynamic stability (measured by transition stability and sequence complexity), we observed that the CAR selectively improved neural resource allocation toward emotional processing, as indicated by a decrease in overall transition stability. And based on entropy analysis, another metric to quantify system’s energy landscape, we observed an increased entropy during emotional condition when suppressed CAR, further validating hypothesis that CAR induced a higher resource allocation toward emotional processing. This allocation could originate from the interplay between CAR induced tonic tuning and task-driven phasic neurotransmitters such as NE, which directly projects into distributed brain networks and serves as a catalyst to regulate internal neural resource in support of ever-changing environmental and cognitive demands (Servan-Schreiber et al. 1990, Young et al. 1998, Clow et al. 2010). For affective processing, glucocorticoids induced by the CAR could interplay with emotional-induced catecholaminergic actions, leading to higher glucose utilization which preferentially support affection (Foote et al. 1987, Hermans et al. 2014). In contrast, an antagonistic effect during high-load working memory could attribute to a reduced energy support due to the interplay between glucocorticoids and norepinephrine, the latter of which may exhibit an inhibitory effect toward executive-control networks during stressful context (Birnbaum et al. 1999) Notably, our observed effect above could be further supported by the fact that a positive correlation between individual differences in CAR (i.e., AUCi) and system stability during emotional discrimination task (suppressed CAR linking to increased transition frequency during emotion compared with neutral). In contrast, we observed an opposite correlation during working memory (suppressed CAR linking to decreased transition frequency during high-load WM compared with low-load). Such antagonistic effect during high-load working memory could attribute to a reduced energy support due to the interplay between glucocorticoids and norepinephrine, the latter of which may exhibit an inhibitory effect toward executive-control networks during stressful context (Birnbaum et al. 1999).

Such opposite energy shifts across emotion and executive functions further indicates a reallocation of neural resource. Researches from both allostasis (Goldstein et al. 2002, McEwen et al. 2020, Bobba-Alves et al. 2022) have pointed out that such reallocation of neurocognitive resource is essential for neurobiological systems which require the ability to anticipate upcoming challenges and stress, and flexibly regulate a limited amount of energy to preferentially support nuanced affective and cognitive functions. Thus, our observed opposite system stability shifts during emotional and executive functioning have important implications into the CAR-mediated proactive effect, through actively modulating neural resource allocation for cognitive and behavioral flexibility, thereby serving as a crucial role in supporting system allostasis. (McEwen 2000, McEwen et al. 2020).

Our study has several limitations. First, our study focused on CAR in males attempting to avoid confounds by menstrual cycles in female, which may impede the generalizability in female populations. Future studies should consider test the proactive effect of CAR in broader populations to gain better generalizability. Second, our study did not directly measure coinciding phasic activity of dopamine and noradrenaline that often interplay with CAR. Future studies should consider implementing in-silicon simulations or techniques to probe dynamic activity of neuromodulators in-vivo to directly model how cortisol interacts with phasic neuromodulators to modulate brain cognitive and affective functions.

**In conclusion**, our study demonstrates the proactive effects of CAR on dynamic reconfigurations of large-scale functional brain networks across emotional and executive functions. Our findings establish a causal link of how CAR could proactively and selectively modulate brain state dynamics via regulating occurrence and overall transition dynamicity of the brain network systems, suggesting a neurocognitive model underlying flexible reallocation of neural resources across affective and cognitive task demands. Our findings provide novel insights into CAR-mediated proactive effects that may serve as a key role in human allostasis to enable brain preparedness for upcoming challenges.

## Supplementary materials

**Figure S1.**
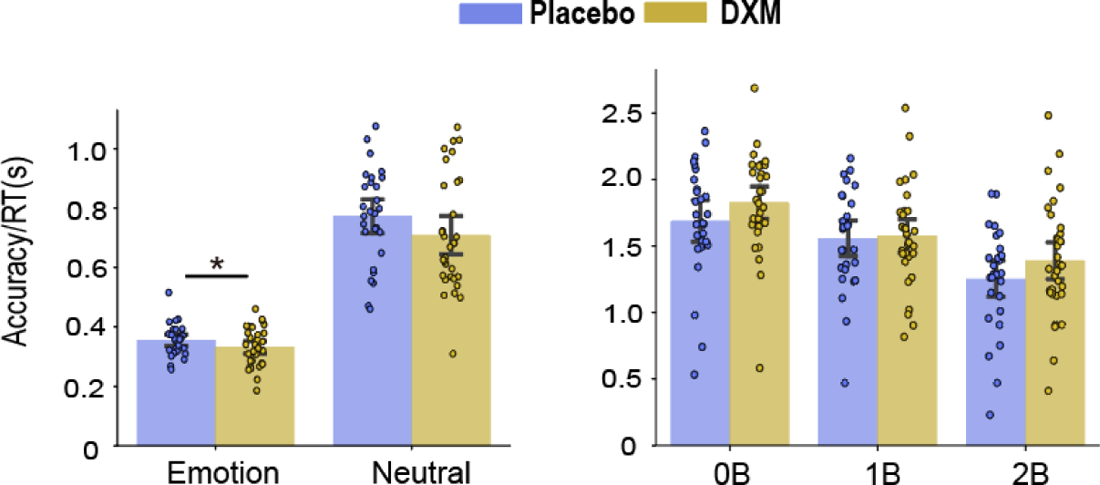
Behavior performance for emotion matching and working memory. A. Bar graphs depict accuracy and accuracy divided by response time (s) during task condition in two groups. Results depict significant main effects of WM-loads for both measures. Notes: *, p<0.05, **, p<0.01. We conducted similar ANOVA for accuracy divide RT and found main effect of only emotion condition (F_1,59_ = 395.85, p <0.001) and no interaction effect (F_1,59_ = 1.04, p = 0.31). Post-hoc t-tests revealed significant higher accuracy divide RT of emotion condition in placebo group compared with that in DXM group (t_59_ = 2.46, p = 0.017) but no significant difference in neutral condition (t_59_ = 1.45, p = 0.15). We conducted similar ANOVA for accuracy divide RT and found significant main effect of workload (F_1,59_ = 66.21, p <0.001).

**Figure S2.**
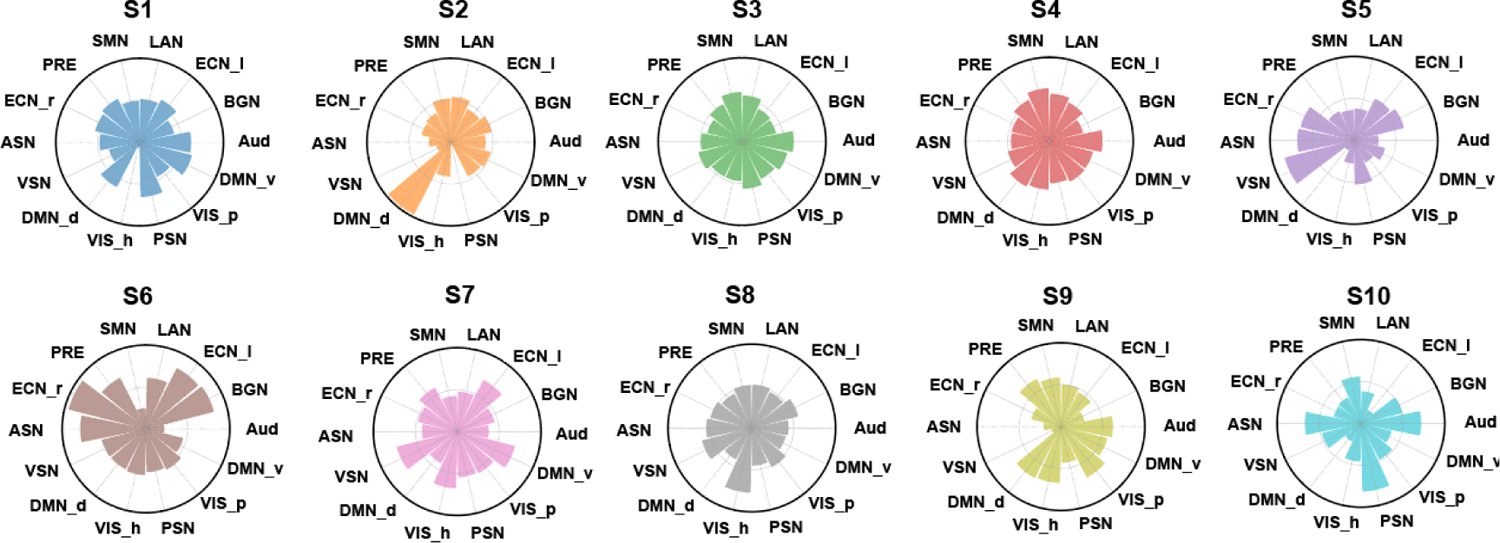
Characterization of hmm inferred brain state. Polar graphs represent each inferred brain state featured by their relative load of 14 canonical networks. The value of each bar indicates the relative activity magnitude compared with the average activity inferred from HMM modeling.

**Figure S3.**
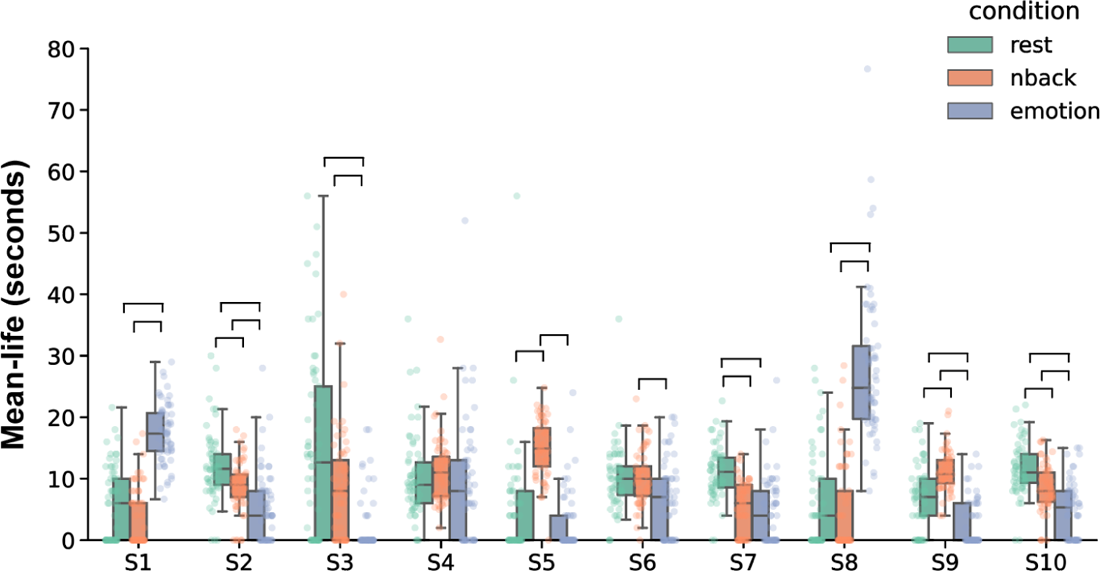
Mean life time for brain state under task conditions Relative mean-life time for each brain state during rest, emotion matching and N-back. Horizontal lines depict significant difference in averaged mean-life time between paired tasks. Notes: horizontal lines represent significant differences in mean life time (p < 0.001, FDR corrected) among three task conditions for each state.

**Figure S4.**
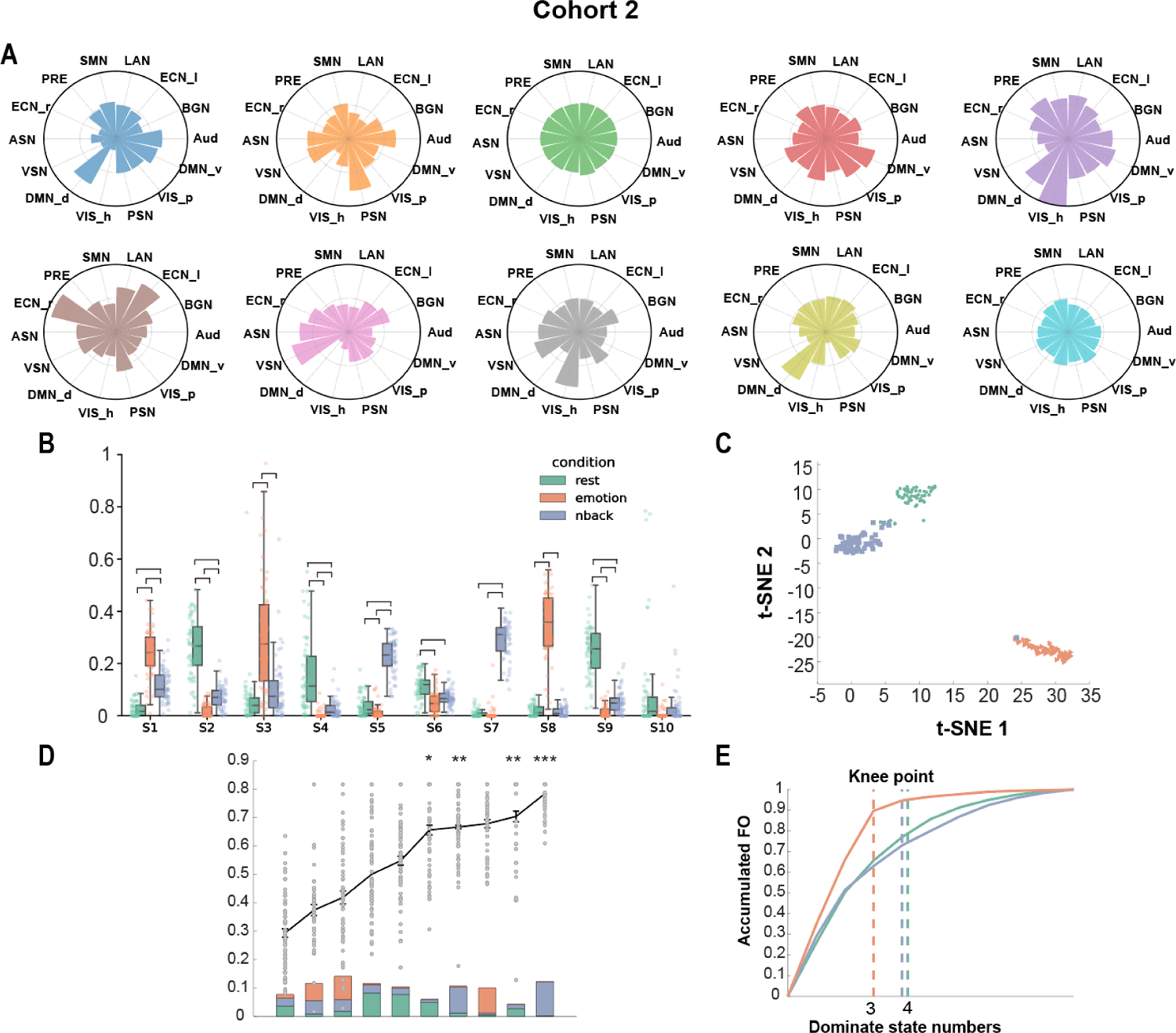
Replicable brain state dynamics in Cohort 2. **(A)** The relative load of 14 canonical networks for each inferred brain state. **(B)**. Relative occupancy rate of each brain state during rest, emotion condition and N-back. **(C)** t-SNE embedding for distribution of occupancy rates of 3 task conditions. **(D)** State specificity across 3 tasks. Line plot depicts specificity index for each state. Dots represent specificity for each state individually. Colored bar depicted relative fractional occupancy for each state while colored portions in each bar represent relative occupancy ratio for that condition. Stats are sorted by their specificity index. **(E)** Dominate state selection based on knee point analysis, vertical dashed lines represent corresponding dominate state numbers determined by knee point for each task separately. Notes: *p_corr_<0.05, **p_corr_<0.01, ***p_corr_<0.001

**Figure S5.**
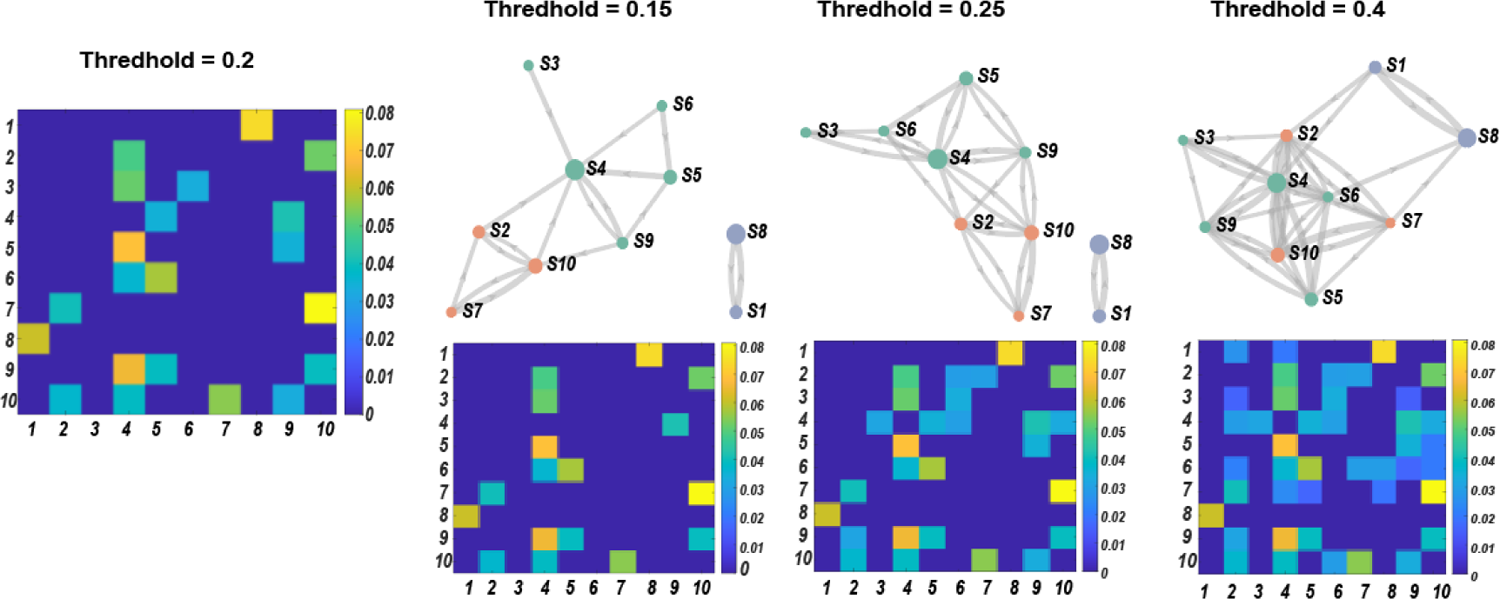
Transition matrix and community detection detections under various threshold. Group transition matrix derived from model and we implemented binary process at various threshold before conducting community detection (0.15,0.2,0.25,0.4). Results showed consistently community assignment patterns regardless of threshold parameter setting.

**Figure S6.**
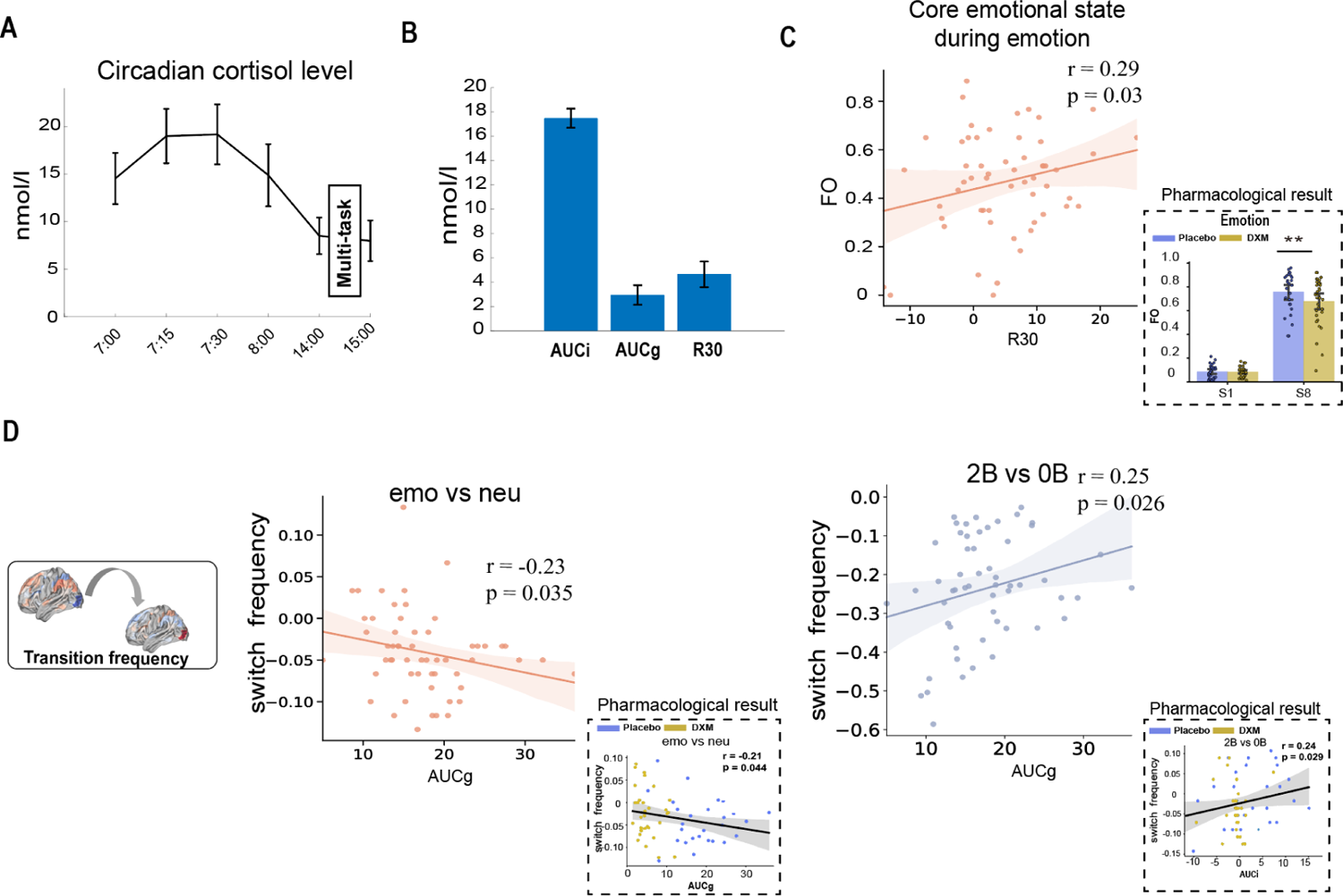
Replicable relationship between individual CAR and brain state dynamics. **(A)** Salivary samplings across 6 time points in Cohort 2. **(B)** AUCi, AUCg and R30 measurement for cortisol awakening response level. **(C)** Similar affective-active state found in Cohort 2 during emotion blocks are positively linked with R30 level. **(D)** Left: Switching frequency shifts from neutral to emotion are negatively correlated with cortisol awakening response (AUCg). Right: switching frequency shifts from 0-back to 2-back are positively correlated with cortisol awakening response (AUCg).

## Reference

Arnsten, A. F. T. (2009). “Stress signalling pathways that impair prefrontal cortex structure and function.” Nature Reviews Neuroscience 10(6): 410–422.

Barulli, D., et al. (2013). “Efficiency, capacity, compensation, maintenance, plasticity: emerging concepts in cognitive reserve.” Trends in Cognitive Sciences 17(10): 502–509.

Bassett, D. S., et al. (2006). “Small-World Brain Networks.” The Neuroscientist 12(6): 512–523.

Bijanzadeh, M., et al. (2022). “Decoding naturalistic affective behaviour from spectro-spatial features in multiday human iEEG.” Nature Human Behaviour.

Birnbaum, S., et al. (1999). “A role for norepinephrine in stress-induced cognitive deficits: α-1-adrenoceptor mediation in the prefrontal cortex.” Biological Psychiatry 46(9): 1266–1274.

Bobba-Alves, N., et al. (2022). “The energetic cost of allostasis and allostatic load.” Psychoneuroendocrinology 146: 105951.

Bullmore, E., et al. (2009). “Complex brain networks: graph theoretical analysis of structural and functional systems.” Nature Reviews Neuroscience 10(3): 186–198.

Capouskova, K., et al. (2022). “Modes of cognition: Evidence from metastable brain dynamics.” Neuroimage 260: 119489.

Chang, C., et al. (2010). “Time-frequency dynamics of resting-state brain connectivity measured with fMRI.” Neuroimage 50(1): 81–98.

Chen, K., et al. (2022). “Hidden Markov Modeling Reveals Prolonged “Baseline” State and Shortened Antagonistic State across the Adult Lifespan.” Cereb Cortex 32(2): 439–453.

Clawson, W., et al. (2019). “Computing hubs in the hippocampus and cortex.” Science Advances 5(6): eaax4843.

Clow, A., et al. (2010). “The cortisol awakening response: More than a measure of HPA axis function.” Neuroscience & Biobehavioral Reviews 35(1): 97–103.

Cornblath, E. J., et al. (2020). “Temporal sequences of brain activity at rest are constrained by white matter structure and modulated by cognitive demands.” Communications Biology 3(1): 261.

Crutchfield, J. P. (2012). “Between order and chaos.” Nature Physics 8(1): 17–24.

de Kloet, E. R., et al. (2019). “Top-down and bottom-up control of stress-coping.” Journal of Neuroendocrinology 31(3): e12675.

Devilbiss, D. M., et al. (2012). “Stress-induced impairment of a working memory task: role of spiking rate and spiking history predicted discharge.”

Ezaki, T., et al. (2017). “Energy landscape analysis of neuroimaging data.” Philosophical Transactions of the Royal Society A: Mathematical, Physical and Engineering Sciences 375(2096): 20160287.

Foote, S. L., et al. (1987). “Extrathalamic modulation of cortical function.” Annual Review of Neuroscience 10(1): 67–95.

Fornito, A., et al. (2012). “Competitive and cooperative dynamics of large-scale brain functional networks supporting recollection.” Proc Natl Acad Sci U S A 109(31): 12788–12793.

Fries, E., et al. (2009). “The cortisol awakening response (CAR): Facts and future directions.” International Journal of Psychophysiology 72(1): 67–73.

Goldstein, D. S., et al. (2002). “Allostasis, homeostats, and the nature of stress.” Stress 5(1): 55–58.

Gu, S., et al. (2017). “Optimal trajectories of brain state transitions.” Neuroimage 148: 305–317.

He, Y., et al. (2023). “Development of brain-state dynamics involved in working memory.” Cerebral Cortex.

Hermans, E. J., et al. (2014). “Dynamic adaptation of large-scale brain networks in response to acute stressors.” Trends in Neurosciences 37(6): 304–314.

Hutchison, R. M., et al. (2013). “Dynamic functional connectivity: promise, issues, and interpretations.” Neuroimage 80: 360–378.

Joëls, M., et al. (2009). “The neuro-symphony of stress.” Nature Reviews Neuroscience 10(6): 459–466.

Joëls, M., et al. (1992). “Control of neuronal excitability by corticosteroid hormones.” Trends in Neurosciences 15(1): 25–30.

Kottaram, A., et al. (2019). “Brain network dynamics in schizophrenia: Reduced dynamism of the default mode network.” Human Brain Mapping 40(7): 2212–2228.

Kunz-Ebrecht, S. R., et al. (2004). “Differences in cortisol awakening response on work days and weekends in women and men from the Whitehall II cohort.” Psychoneuroendocrinology 29(4): 516–528.

Lamont, E. W., et al. (1998). “Infusion of the dopamine D1 receptor antagonist SCH 23390 into the amygdala blocks fear expression in a potentiated startle paradigm.” Brain research 795(1-2): 128–136.

Law, R., et al. (2020). Chapter Eight - Stress, the cortisol awakening response and cognitive function. International Review of Neurobiology. A. Clow and N. Smyth, Academic Press. 150: 187–217.

McEwen, B. S. (2000). “Allostasis and Allostatic Load: Implications for Neuropsychopharmacology.” Neuropsychopharmacology 22(2): 108–124.

McEwen, B. S., et al. (2020). “Revisiting the Stress Concept: Implications for Affective Disorders.” The Journal of Neuroscience 40(1): 12–21.

McReynolds, J. R., et al. (2010). “Memory-enhancing corticosterone treatment increases amygdala norepinephrine and Arc protein expression in hippocampal synaptic fractions.” Neurobiology of Learning and Memory 93(3): 312–321.

Meer, J. N. v. d., et al. (2020). “Movie viewing elicits rich and reliable brain state dynamics.” Nature Communications 11(1): 5004.

Menon, V., et al. (2010). “Saliency, switching, attention and control: a network model of insula function.” Brain Struct Funct 214(5-6): 655–667.

Moriarty, A. S., et al. (2014). “Cortisol awakening response and spatial working memory in man: a U-shaped relationship.” Human Psychopharmacology: Clinical and Experimental 29(3): 295–298.

Munn, B. R., et al. (2021). “The ascending arousal system shapes neural dynamics to mediate awareness of cognitive states.” Nature Communications 12(1): 6016.

Presigny, C., et al. (2022). “Colloquium: Multiscale modeling of brain network organization.” Reviews of Modern Physics 94(3): 031002.

Pruessner, J. C., et al. (1997). “Free Cortisol Levels after Awakening: A Reliable Biological Marker for the Assessment of Adrenocortical Activity.” Life Sciences 61(26): 2539–2549.

Ray, K. L., et al. (2020). “Dynamic reorganization of the frontal parietal network during cognitive control and episodic memory.” Cognitive, Affective, & Behavioral Neuroscience 20(1): 76–90.

Roozendaal, B., et al. (2009). “Stress, memory and the amygdala.” Nature Reviews Neuroscience 10(6): 423–433.

Satopaa, V., et al. (2011). Finding a “Kneedle” in a Haystack: Detecting Knee Points in System Behavior. 2011 31st International Conference on Distributed Computing Systems Workshops.

Seeley, W. W., et al. (2007). “Dissociable Intrinsic Connectivity Networks for Salience Processing and Executive Control.” The Journal of Neuroscience 27(9): 2349.

Servan-Schreiber, D., et al. (1990). “A Network Model of Catecholamine Effects: Gain, Signal-to-Noise Ratio, and Behavior.” Science 249(4971): 892–895.

Shine, J. M. (2019). “Neuromodulatory Influences on Integration and Segregation in the Brain.” Trends in Cognitive Sciences 23(7): 572–583.

Shirer, W. R., et al. (2012). “Decoding subject-driven cognitive states with whole-brain connectivity patterns.” Cereb Cortex 22(1): 158–165.

Steptoe, A., et al. (2016). Cortisol awakening response. Stress: Concepts, cognition, emotion, and behavior, Elsevier: 277–283.

Stevner, A. B. A., et al. (2019). “Discovery of key whole-brain transitions and dynamics during human wakefulness and non-REM sleep.” Nature Communications 10(1): 1035.

Telesford, Q. K., et al. (2016). “Detection of functional brain network reconfiguration during task-driven cognitive states.” Neuroimage 142: 198–210.

Van der Maaten, L., et al. (2008). “Visualizing data using t-SNE.” Journal of machine learning research 9(11).

van Stegeren, A. H., et al. (2007). “Endogenous cortisol level interacts with noradrenergic activation in the human amygdala.” Neurobiology of Learning and Memory 87(1): 57–66.

Vidaurre, D., et al. (2017). “Brain network dynamics are hierarchically organized in time.” Proceedings of the National Academy of Sciences 114(48): 12827.

Xin, F., et al. (2018). “Oxytocin Modulates the Intrinsic Dynamics Between Attention-Related Large-Scale Networks.” Cerebral Cortex 31(3): 1848–1860.

Xiong, B., et al. (2021). “Brain preparedness: The proactive role of the cortisol awakening response in hippocampal-prefrontal functional interactions.” Progress in Neurobiology 205: 102127.

Yarkoni, T., et al. (2011). “Large-scale automated synthesis of human functional neuroimaging data.” Nature Methods 8(8): 665–670.

Young, C. B., et al. (2017). “Dynamic Shifts in Large-Scale Brain Network Balance As a Function of Arousal.” The Journal of Neuroscience 37(2): 281–290.

Young, E. A., et al. (1998). “The Role of Mineralocorticoid Receptors in Hypothalamic-Pituitary-Adrenal Axis Regulation in Humans1.” The Journal of Clinical Endocrinology & Metabolism 83(9): 3339–3345.

